# Large Data on the Small Brain: Population-wide Cerebellar Growth Models of Children and Adolescents

**DOI:** 10.1101/2023.04.26.538263

**Authors:** Carolin Gaiser, Rick van der Vliet, Augustijn A. A. de Boer, Opher Donchin, Pierre Berthet, Gabriel A. Devenyi, M. Mallar Chakravarty, Jörn Diedrichsen, Andre F. Marquand, Maarten A. Frens, Ryan L. Muetzel

## Abstract

In the past, the cerebellum has been best known for its crucial role in motor function. However, increasingly more findings highlight the importance of cerebellar contributions in cognitive functions and neurodevelopment. Using large scale, population-wide neuroimaging data, we describe and provide detailed, openly available models of cerebellar development in childhood and adolescence, an important time period for brain development and onset of neuropsychiatric disorders. Next to a traditionally used anatomical parcellation of the cerebellum, we generated growth models based on a recently proposed functional parcellation. In both, we find an anterior-posterior growth gradient mirroring the age-related improvements of underlying behavior and function, which is analogous to cerebral maturation patterns and offers new evidence for directly related cerebello-cortical developmental trajectories. Finally, we illustrate how the current approach can be used to detect cerebellar abnormalities in clinical samples.

## 2. Introduction

The cerebellum is known to be engaged in a broad spectrum of functions. While its involvement in motor control is best documented, recent efforts have made clear that it is also involved in cognitive function. Given that the cerebellum is strongly interconnected with the cerebral cortex, with cerebellar functional subunits being involved in a wide array of motor and cognitive tasks (Buckner et al., 2011; Strick et al., 2009), these recent findings come as no surprise. Yet, despite converging evidence on the importance of the cerebellum for brain function, limited work has explored how the cerebellum develops through childhood and adolescence.

The cerebellum is one of the first structures in the brain to start cellular differentiation, with a rapid growth period in the third trimester of pregnancy and in the first postnatal year, but it is one of the last to complete maturity (Limperopoulos et al., 2005; Wang & Zoghbi, 2001). Given this protracted developmental time course, it is especially vulnerable to genetic and environmental stressors disrupting development (Wang et al., 2014; Wang & Zoghbi, 2001). It thus could be a key node in various neurodevelopmental disorders and has the potential to serve as a crucial biomarker.

In addition to overlooking its role in higher cognitive function in the past, challenges in *in vivo* imaging of the cerebellum have likely hampered the study of this important structure. The anatomical location of the cerebellum and its tightly folded cortex have made it a more challenging structure to image since the acquisition field of view and head coils are often optimized for imaging the cerebrum. Additionally, high resolution images are needed for precise segmentations as well as anatomical and functional mapping. The adoption of high magnetic field strengths of 3T and beyond in tandem with the development of dedicated automated segmentation and parcellation tools (Diedrichsen et al., 2009; Faber et al., 2022; Han et al., 2020; Park et al., 2014) has made the analysis of cerebellar imaging data more accessible.

As childhood and adolescence represent a time of increased risk for psychiatric and developmental problems (Paus et al., 2008), it is crucial to improve our understanding of cerebellar development during this period. For this reason, robust and detailed reference models of neurodevelopmental trajectories are needed, which recently has become a thriving area of research (Bethlehem et al., 2022; Rutherford et al., 2022b). Normative modeling of brain imaging data is particularly well suited to this task and provides an analysis framework that is able to model biological heterogeneity at the level of the individual while also accommodating site effects (Kia et al., 2022; Rutherford et al., 2022a). This new framework allows for tracking the development of a given individual against expected centiles in variation of a reference model, without needing to assume that clinical populations are homogeneous, analogous to growth charts in pediatric medicine. In the context of psychopathology, this approach has recently shown to increase sensitivity and to better characterize inter-individual heterogeneity in regional brain volumes compared to case-control studies (Remiszewski et al., 2022; Wolfers et al., 2018). By embedding normative modeling of imaging data within a federated learning framework, sharing of such models becomes possible without data privacy concerns. This not only means that smaller datasets can benefit from informative hyperpriors of a reference model based on much larger datasets, but also that models can be adapted and updated as more data becomes available (Kia et al., 2022). Using this framework to establish normative models of the cerebellum will therefore prove extremely useful to detect deviations in cerebellar development on the level of individuals and to map these deviations to behavioral and clinical phenotypes.

Traditionally, the cerebellum has been subdivided in the medial-to-lateral direction into vermis and hemispheres, and in the anterior-to-posterior direction into lobules. However, more recently, *functional magnetic resonance imaging* (fMRI) has shown that functional boundaries in the cerebellum do not align with classical anatomical subdivisions (King et al., 2019). Instead, an alternative functional parcellation containing at least 10 regions has been identified, which corresponds well to earlier proposed cerebro-cerebellar network parcellations (Buckner et al., 2011), and that are characterized by the motor and cognitive features that elicit activity in the parcels (King et al., 2019).

In the current study, we describe and provide openly available normative models of anatomical and functional subregions of the cerebellum from a large pediatric population that 1) can be used as reference models to obtain accurate normative ranges, also in smaller datasets, by benefitting from informative hyperpriors based on a large sample, and that 2) can be updated with data from new sites and extended age ranges without the necessity of sharing sensitive patient or participant data. We furthermore illustrate the usefulness and practicality of the current approach by mapping the deviations from typical cerebellar development at the level of the individual in a subpopulation of children with autistic traits (Constantino et al., 2003). These models have the potential to facilitate and maximize the use of cerebellar outcomes in neuroimaging research and as a result aid to better understand the role of the cerebellum in typical as well as atypical neurodevelopment.

## 3. Methods

### 3.1. Participants

Participants were part of the Generation R Study, a population-based, prospective cohort study from fetal life onward (Kooijman et al., 2016). Between 2002 and 2006, 9,778 pregnant women living in Rotterdam, The Netherlands were enrolled in the study. Data from the children and caregivers were collected at several time points. In total, MRI data from 5,185 unique individuals were obtained across three time points. 1,070 participants (mean age = 7.9) visited during the first assessment, 3,992 participants (mean age = 10.2) during the second, and 3,725 participants (mean age = 14.0) visited the testing center at the third assessment. After exclusion of participants with incomplete T_1_-weighted scans (N = 1,214), scans without complete consent form (N = 122), scans with incidental findings (N = 73), and scans with low image quality ratings (N = 454), a total of 7,270 scans from 4,862 individuals were available for statistical analysis. Written informed consent and/or assent was obtained from all participants, and the study was approved by the Medical Ethical Committee of the Erasmus Medical Center.

### 3.2. Neuroimaging acquisition

Scans were acquired using two different MRI scanners. In the first wave, data were collected on a GE MR750 Discovery system, and data from all subsequent assessments were collected on a study-dedicated GE MR750w system (General Electric Healthcare, Wisconsin, USA). High resolution T_1_-weighted MRI scans were acquired using an inversion recovery fast spoiled gradient recalled sequence (IR-FSPGR) using the following parameters: Wave 1: T_R_=10.3 ms, T_E_=4.2 ms, T_I_=350 ms, flip angle = 16°, acquisition time = 5 min 40 s, field of view = 230.4 x 230.4 mm, 0.9x0.9x0.9 mm^3^ isotropic resolution. Wave 2 and 3: T_R_=8.77 ms, T_E_=3.4 ms, T_I_=600 ms, flip angle = 10°, acquisition time = 5 min 20 s, field of view = 220 x 220 mm, 1x1x1 mm^3^ isotropic resolution.

### 3.3. Image analysis

#### 3.3.1. Pre-processing

Images from the first measurement wave were resampled to 1 mm isotropic resolution to match data from the second and third assessments. Images were then pre-processed using the SMRIPrep tool (Esteban et al., 2021). Briefly, non-brain tissue was removed, voxel intensities were adjusted for B_1_ inhomogeneities, and images were then linearly and eventually nonlinearly registered to standard stereotactic space (MNI152 NLin2009cAsym 1x1x1 mm resolution) using ANTs (github: https://github.com/ANTsX/ANTs.git). The tissue segmentation procedure resulted not only in binary classifications of voxels, but also in per-voxel tissue class probability estimates. These probabilities were then interpreted as the grey and white matter and cerebrospinal fluid densities for a given voxel. Further, the nonlinear registration produced a nonlinear warp file (which included the linear initialization) from which we calculated the determinant of the Jacobian matrix for each voxel. This determinant was used as a measure of volume of that voxel relative to its volume in standard stereotactic space.

#### 3.3.2. Anatomical parcellation

The cerebellum was parcellated in the native space into 35 anatomical subdivisions using the MAGeT pipeline (Chakravarty et al., 2013; Park et al., 2014). The MAGeT Brain framework uses an automated segmentation algorithm based on five manually segmented MR images from healthy participants. We used non-linear registration to these five manually segmented images to automatically segment a small number of individual “template” images chosen from the study dataset. In the current study, we selected seven unique and representative images from the three time points, resulting in 21 study-specific template images. Each of the five manually segmented atlases were applied to the 21 study-specific templates, resulting in 105 template segmentations in total. In a final step, each image in the dataset is segmented by non-linear registration to each of the templates resulting in 105 segmentations for each input image (Chakravarty et al., 2013). These were then fused using voxel-wise majority voting to create a final segmentation for each input image. Volumes for each of the 35 anatomical parcellations are generated in mm^3^ by the MAGeT pipeline. *Supplementary Figure 1* shows a representative automatically labelled segmentation from one individual. This computationally intensive approach has been shown to have better test-retest reliability than other segmentation techniques (Park et al., 2014).

#### 3.3.3. Functional parcellation

As lobular boundaries of the cerebellum have shown limited correspondence with functional boundaries, we also employed the functional subregions proposed by King and colleagues (King et al., 2019). Ten functional regions of the cerebellum were identified using fMRI data from a large multi-domain task battery (MDTB) and labelled according to the cognitive features that best described the task conditions (*1: Left-hand (motor) presses, 2: Right-hand (motor) presses, 3: Saccades, 4: Action observation, 5: Divided attention (left hemisphere), 6: Divided attention (right hemisphere), 7: Narrative, 8: Word comprehension, 9: Verbal fluency, 10: Autobiographical recall)*. This parcellation was shown to successfully predict functional boundaries in a new set of motor, cognitive, affective, and social tasks, surpassing existing task-free and anatomical parcellations (King et al., 2019). We used the MNI-aligned version of the MDTB atlas. Mean *grey matter density* (GMD) and *white matter density* (WMD) (see 3.3.1, values closer to 1 indicate a high probability of a given tissue type in that voxel) and volumes (defined as sum of the Jacobian determinants) were extracted for each of the ten functional parcellations (see 3.3.1 Pre-processing).

#### 3.3.4. Image Quality Control

To ensure segmentation quality, anatomical segmentations were visually inspected by two expert raters (C.G., N.D.). A custom-made MATLAB app (version R2021b, Mathworks, USA) was used to inspect PNG files of all slices of each scan, and the segmentation quality was rated on a 3-point scale (*‘’Good”, “Sufficient”, “Bad”)* based on inaccuracies in the parcellation, complete coverage of the cerebellum, and motion or other artifacts. Scans rated as *“Bad”* (i.e., cases without full coverage of the cerebellum, scans with substantial artifacts, and/or scans with marked inaccuracies in the parcellation) were subsequently excluded from further analyses. A subset of 600 scans were inspected by both raters to assess *inter-rater reliability* (IRR).

### 3.4. Normative models

To generate normative models for anatomical and functional subregions of the cerebellum, we made use of the PCNtoolkit python package version 0.27 (de Boer et al., 2022; Marquand et al., 2016; Rutherford et al., 2022b) using Python 3.10.6. The PCNtoolkit allows for 1) obtaining normative ranges (Marquand et al., 2016), 2) modeling individual heterogeneity to uncover clinically significant deviations (Marquand et al., 2016; Remiszewski et al., 2022; Rutherford et al., 2022a), 3) correcting for batch-effects, such as differences between scanners (Kia et al., 2022), and 4) enabling data sharing without disclosing sensitive patient or participant information (Kia et al., 2022). The *Hierarchical Bayesian Regression* (HBR) approach implemented in PCNtoolkit solves these problems by using shared priors from which the site-specific parameters and hyperparameters can be learned, and by providing a framework in which generated hyperparameters from previous analyses can be made available to new sites without the original data (i.e., a federated approach). Previously, we have shown that the HBR approach can be used to perform meaningful inferences across longitudinal time points, even when subsequent waves are scanned on different scanners (Gaiser et al., 2023).

Using HBR, we estimated normative models of the cerebellum using both the volumes from the anatomical parcellation and the morphological indicators (i.e., grey and white matter densities as well as volumes) of the functional parcellation from age, for each *region of interest* (ROI) separately. Sex and scanner were modeled as batch-effects. A graphical representation of the normative model and its parameters can be found in *Supplementary Figure 2*. For both the anatomical and functional parcellations, we split the dataset into a training set (50%) and test set (50%) using the sex and scanner site variables to ensure equal distribution of sex and both scanners in both sets. We generated linear and 3^rd^ order b-spline models with 5 evenly spaced knot points (all available at: https://pcnportal.dccn.nl/). Model performance of both linear and b-spline models were evaluated using Leave-one-out cross-validation (LOO). We employed a sinh-arcsinh likelihood (SHASHb) to accommodate non-Gaussian distributions (de Boer et al., 2022) and modeled random effects in intercept, slope, and variance (sigma) on the batch-effects (sex and scanner). Inference was performed using Markov chain Monte Carlo methods (see Kia et al., 2022 and de Boer et al., 2022 for full details). Four chains with 2000 samples each were generated. The first 500 samples of each chain were used as tuning samples and were removed from further analysis. Model outputs include the posterior distributions of the parameters and deviations from the normative range (z-score which are free of batch-effects) for each individual in the test set.

### 3.5. Clinical validation of models using Social Responsiveness Scores (SRS)

To illustrate the utility of the cerebellar normative models, we examine whether deviations in cerebellar growth are present in children who are likely to fall on the autism spectrum according to the *Social Responsiveness Scale* (SRS). The SRS has been shown to quantitatively assess subclinical and clinical autistic traits (Constantino et al., 2003). A shortened 18-item version was administered via questionnaire at the age of 8 years (Blanken et al., 2015). For each ROI, we contrast the z-scores between children likely to fall on the autism spectrum (raw score on SRS >= 90^th^ percentile, N = 198) to the remainder of the cohort (N = 2,012; children without SRS information excluded). Previously, it has been shown that using these deviation scores can uncover more precise case-control effects and are able to characterize clinically relevant differences in morphology on an individual level (Remiszewski et al., 2022; Rutherford et al., 2022a). We therefore illustrate the percentage of children with high SRS scores that have a large deviation from the normative range (z > 1.96 / z < −1.96, critical z for 95% confidence interval) per ROI. Using this definition, we expect roughly 2.5% of a typically developing population to have a large negative or a large positive z-score, respectively in almost all ROIs of the anatomical and functional parcellations. Significance of the percentage of children with large deviations at *p* = 0.05 level were evaluated using Binomial testing (observed vs. expected number of participants with z > 1.96 / z < −1.96 in high SRS and typical children, given a null hypothesized probability of *p_0_* = 0.025, one-sided).

### 3.6. Non-response analysis

Given the prospective, longitudinal nature of the study, it is important to understand the impact of loss-to-follow-up. We therefore tested random drop-out in our study by examining possible differences between participants included and excluded in the current analysis in terms of the following descriptive characteristics: sex, parental national origin (Dutch, Non-Dutch but European, Non-European; obtained from birth records), monthly household net income (low = < 1,200€, middle = 1,200€ - 3,200€, high = >3,200€; obtained from questionnaire), maternal education (higher education pursued or not; obtained from questionnaire), IQ and behavioral problems. Nonverbal IQ scores, normalized for sex and age, were measured using 2 subsets (Mosaics [spatial visualization] and Categories [abstract reasoning]) of the *Snijders-Oomen Nonverbal Intelligence Test* (SON-IQ) in the first measurement wave (mean age = 7.9). Behavioral problems were measured using the *Child Behavioral Checklist* (CBCL) (Achenbach, 2001) in the second measurement wave (mean age = 10.1). We dichotomized behavioral problems according to maternal reports (scoring above 80^th^ percentile: behavioral problems present; below 80^th^ percentile: behavioral problems not present).

## 4. Results

### 4.1. Sample characteristics and non-response analysis

**Figure 1** shows the age and scanner distribution of the 7,270 scans, stemming from 4,862 unique individuals, available for statistical analysis. In the first measurement wave, participants included in the analysis had a mean age of 7.9 years (range = [6.1-10.7], N = 974), in the second wave a mean age of 10.1 years (range = [8.6 – 12.0], N = 3,785), and in the third wave a mean age of 14.0 years (range = [12.6 - 17.1], N = 2,511). 2,734 (56.2%) individuals were measured once, 1,848 (38.0%) individuals twice, and scans in all three measurement waves were acquired from 280 (5.8%) individuals. The population was split into training and test sets as described in the methods (see *3.4. Normative models*). An overview of the sample characteristics of the training and test sets is shown in *Supplementary Table 1*. Inspection of the table suggests that random allocation of the subjects to trainings- and test sets resulted in groups that are representative of the population as a whole, also when considered at the sub-group level of measurement wave. A non-response analysis was conducted to examine if the participants included in the training and test datasets differed significantly from those excluded due to missing T_1_-weighted scans, incidental findings, incomplete/revoked consent forms, or poor image quality. Participants included in the dataset were more likely to be female (χ^2^ = 8.38, *p* = 0.004), of Dutch national origin (χ^2^ = 181.10, *p* < 0.001), have higher maternal education (χ^2^ = 27.10, *p* < 0.001), and to have a middle or high household income (χ^2^ = 15.48, *p* < 0.001) and less likely to have an IQ below 85 (χ^2^ = 63.21, *p* < 0.001). Behavioral problems as measured by the CBCL did not differ between excluded and included participants (weighted total problem score; χ^2^ = 2.41, *p* = 0.120).

**Figure 1:**
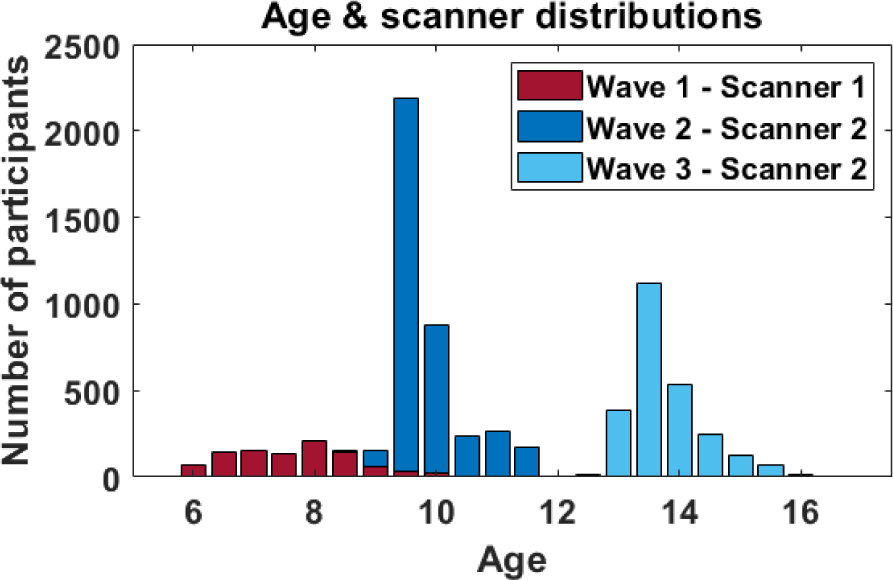
Histogram of age and scanner distributions in the Generation R cohort.

### 4.2. Image quality control: Interrater reliability

After visual inspection of all available scans (N = 8,787 before exclusions), 454 (5.2%) were excluded due to low quality ratings, 593 were rated as *Sufficient* (6.7%), and 7740 as *Good* (88.1%). Inter-rater reliability between both raters (C.G., N.D.) was examined using a subset of 600 double rated scans. Results showed strong agreement on usability (usable or not usable) of the scans between raters (Cohen’s κ = 0.83, CI = [0.72 0.94]).

### 4.3. Normative models of the cerebellum

We modelled the effect of age on cerebellar features of interest (volumes, GMD, WMD) while correcting for batch-effects of sex and scanner (model parameters are illustrated in *Supplementary Figure 2*). Normative models of all previously described cerebellar ROIs were successfully generated. Both linear and b-spline models perform equally well in this age range based on Leave-one-out cross-validation (LOO) (*Supplementary Table 2*). Following the principle of parsimony, the linear models will be described, however b-spline models are made available as well as they might offer more flexible modeling options in future applications. Posterior distributions of all model parameters and ROIs converged well (>95% of 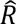 below 1.1; (Brooks & Gelman, 1998)) and were visually inspected using a built-in function of the PCNtoolkit. In the following sections, ROIs of the anatomical and functional parcellations will be described separately.

#### 4.3.1. Anatomical parcellation

We fit a normative model to investigate age-related effects in volume for each of the 35 anatomical ROIs (Park et al., 2014). **Figure 2** illustrates the growth for each ROI using the mean posterior distribution of the age β coefficient (slope). Standardized coefficients are used to ease the comparison between outcomes and results are stratified by sex. As expected for this age range, we see increasing volumes throughout all ROIs. The corpus medullare, the white matter of the cerebellum, shows the most marked increases in volume in both females and males. Interestingly, we see a growth gradient, starting with smaller age-related effects on volume in the anterior cerebellum (Lobules III – VI), and increasingly larger age-related effects in the posterior cerebellum (Lobules VII – IX) with the largest effects, besides the corpus medullare, found in the flocculus (Lobules X). Growth trajectories of example ROIs in the anterior (left Lobule VI) and posterior (left Crus I) cerebellum are shown, as well as for the left corpus medullare (**Figure 2B**). **Figure 2A** furthermore depicts sex differences in age β coefficients (slopes). On average, age-related coefficients were slightly higher for females (mean standardized β across ROIs = 0.178; 95% CI mean = [0.153 0.202]) than for males (mean standardized β across ROIs = 0.149; 95% CI mean = [0.128 0.170]), with larger effects in lobules VIIIA (left hemisphere: difference in standardized β = 0.142; and right hemisphere: difference in standardized β = 0.093), and left lobule X (difference in standardized β = 0.110). An overview of the age β coefficients for all ROIs can be found in *Supplementary Table 3* and growth trajectories of all ROIs stratified by sex in *Supplementary Figure 3A-B*.

**Figure 2:**
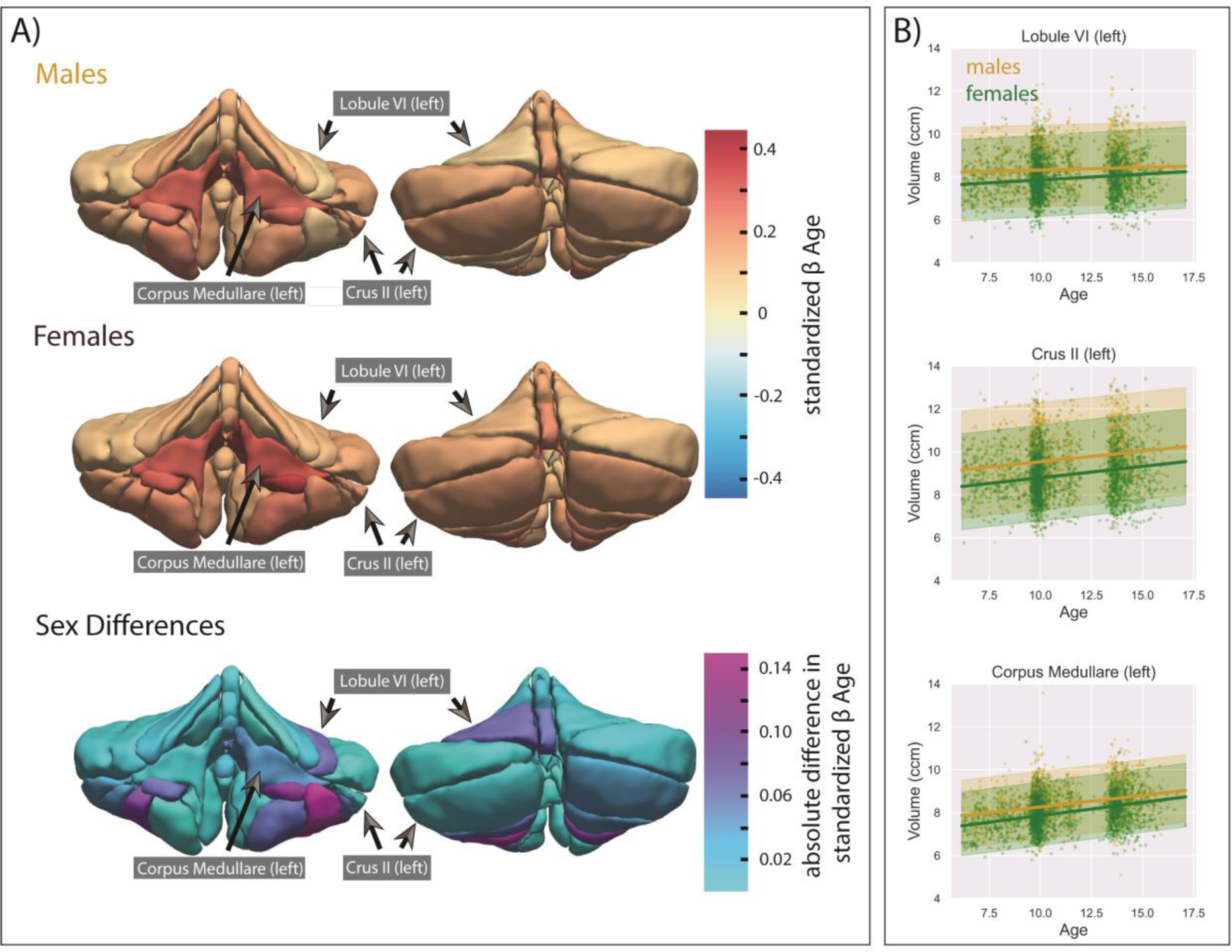
Effect of age on volume in the anatomical parcellation. **A)** Mean posterior distribution for the standardized age β coefficient (slope) for each anatomical ROI and absolute differences in effect sizes (standardized β) between males and females are illustrated. **B)** Trajectories of males (in yellow) and females (in green in 3 example ROIs: left Lobule VI (anterior cerebellum), left Crus II (posterior cerebellum), and left corpus medullare (white matter tract). The bold lines represent the mean trajectories, shaded areas represent what is within 2 standard deviations of the mean.

#### 4.3.2. Functional parcellation

The 10 regions of the MDTB parcellation are illustrated in **Figure 3**. Analogous to the anatomical parcellations, a normative model for the volume, *grey matter density* (GMD), and *white matter density* (WMD) was fit for each ROI of the functional parcellation. In **Figure 4**, we visualize the developmental trajectories using again the mean posterior distribution of the standardized age β coefficient (slope). Results are shown stratified by sex. Like previously seen in the anatomical parcellations, increases in volume are evident throughout all functional parcellations in males and females in this age range (**Figure 4A I & IV**). Once more, smaller age-related effects in volumes are present in the anterior parcels, known to be related to motor behavior, compared to posterior cerebellar regions, which comprise parcels associated with a range of cognitive processes. This trend can be seen even more strikingly in the GMD (**Figure 4A II & V**) and WMD (**Figure 4A III & VI**) models. While it is well-documented that GMD decreases and WMD increases in the brain during this age range, we again see a clear distinction between anterior motor regions and posterior cognitive regions. This is further illustrated by the growth trajectories of an example anterior motor (*1: Left hand presses)* and posterior cognitive (*5: Divided attention (left))* ROI. Steeper slopes, and thus more developmental changes, are observed in posterior cognitive regions compared to anterior motor regions (**Figure 4BI,II & III**). Interestingly, slight differences in sex can be observed with slower changes in GMD and WMD in females compared to males, particularly in the right hemisphere (mean standardized β across ROIs [95% CI mean]: volume males = 0.277 [0.251 0.304], volume females = 0.289 [0.256 0.322], GMD males = −0.138 [−0.206 −0.071], GMD females = −0.056 [−0.124 0.012], WMD males = 0.256 [0.165 0.347], WMD females = 0.184 [0.098 0.270]). Sex differences per ROI are illustrated in **Figure 4A VII, VIII & IX**). As with the anatomical ROIs, an overview of the age β coefficients for all functional ROIs can be found in *Supplementary Table 4* and growth trajectories stratified by sex in *Supplementary Figure 3C-D*.

**Figure 3:**
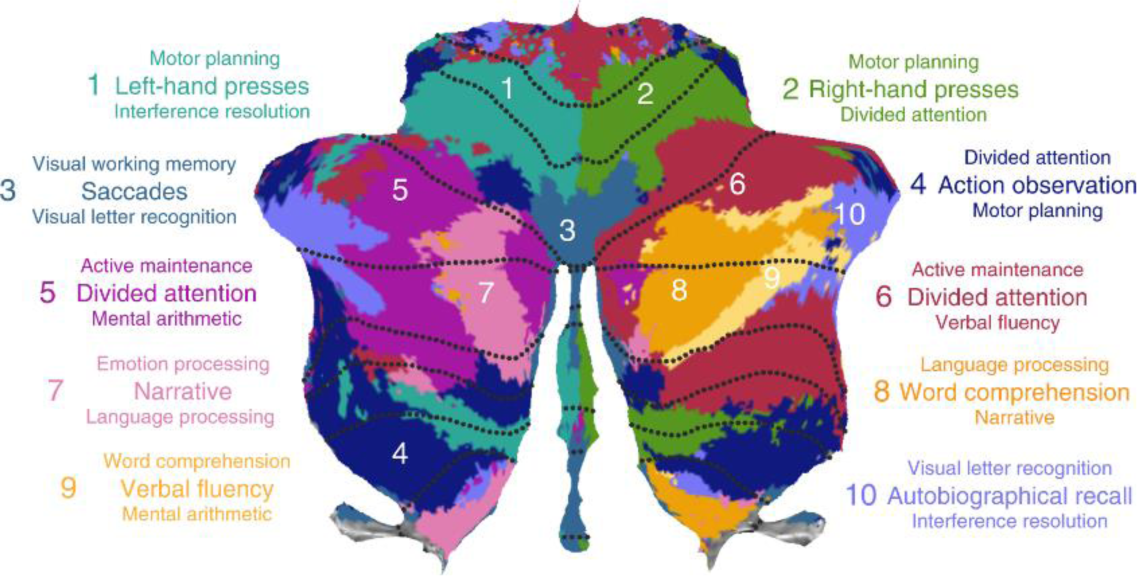
MDTB functional atlas regions. Figure from King et al. 2019.

**Figure 4:**
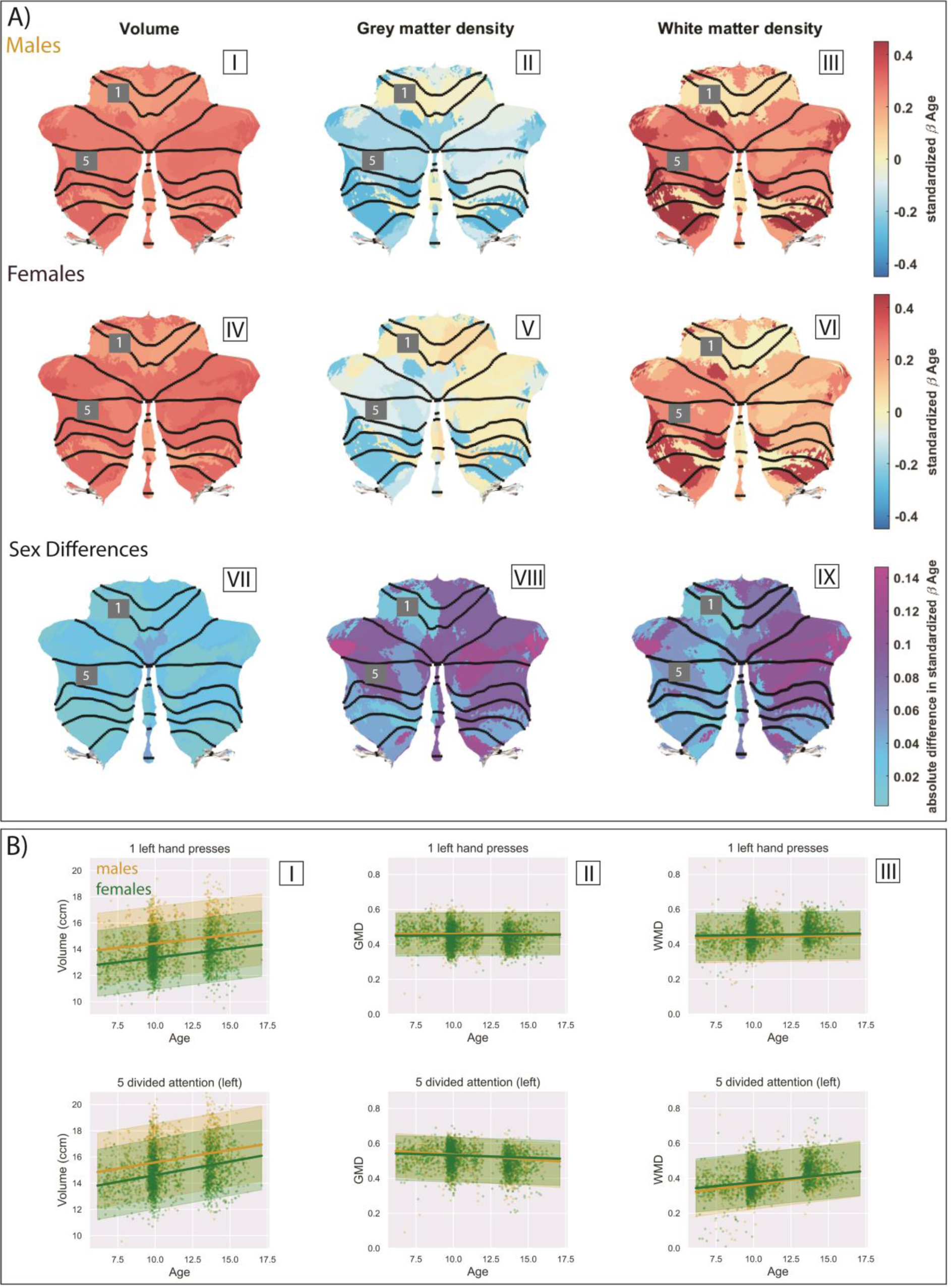
Effect of age on volume in the anatomical parcellation. **A)** Mean posterior distribution for the standardized age β coefficient (slope) for each functional ROI of the MDTB atlas (I-VI) and absolute differences in effect size (standardized β) between males and females are illustrated (VII-IX). **B)** Trajectories of males (in yellow) and females (in green) for 2 example ROIs. 1: *Left hand presses* (anterior cerebellum) and 5: *Divided attention (left)* (posterior cerebellum). The bold lines represent the mean trajectories, shaded areas represent what is within 2 standard deviations of the mean.

### 4.4. Relating large deviations from the normative model to clinical or behavioral phenotypes

To further highlight the utility of the normative models, we compare the normative estimates in individuals with high levels of autistic traits (high SRS scores) to typically developing participants (having low SRS scores). Specifically, we characterize data as having large deviations in the normative model if the value of their normative estimate was > 1.96 or smaller −1.96 (i.e., upper and lower tails of the distribution). As expected using this definition, we observed roughly 2.5% of typically developing participants had a large negative or a large positive z-score, respectively in almost all ROIs of the anatomical and functional parcellation (**Figure 5**, *Supplementary Figure 4*). However, for participants with autistic traits (high SRS) this was not the case as more individuals than expected were represented in the extreme ends of the distribution. In the anatomical parcellation, a higher percentage of participants with high SRS scores presented with large negative z-scores (smaller volume than expected) throughout various ROIs (**Figure 5A**), specifically in vermal and lobular regions of the anterior and superior posterior cerebellum (significant percentage with large deviations binomial test at p<0.05: Crus I (left), VIIIB (left), vermal region III, Lobule VI (right)). Large positive deviations (larger volumes than expected) can be observed in typical participants with low SRS (significant percentage with large deviations binomial test at p<0.05: Left Lobules VI and Crus I, vermal region VIIA, and right Lobule VIIIB). In participants with high SRS scores large positive deviations seem to be less prevalent overall but significant deviations can be indeed observed in the left Lobule IV and vermal region IX (*p*<0.05).

**Figure 5:**
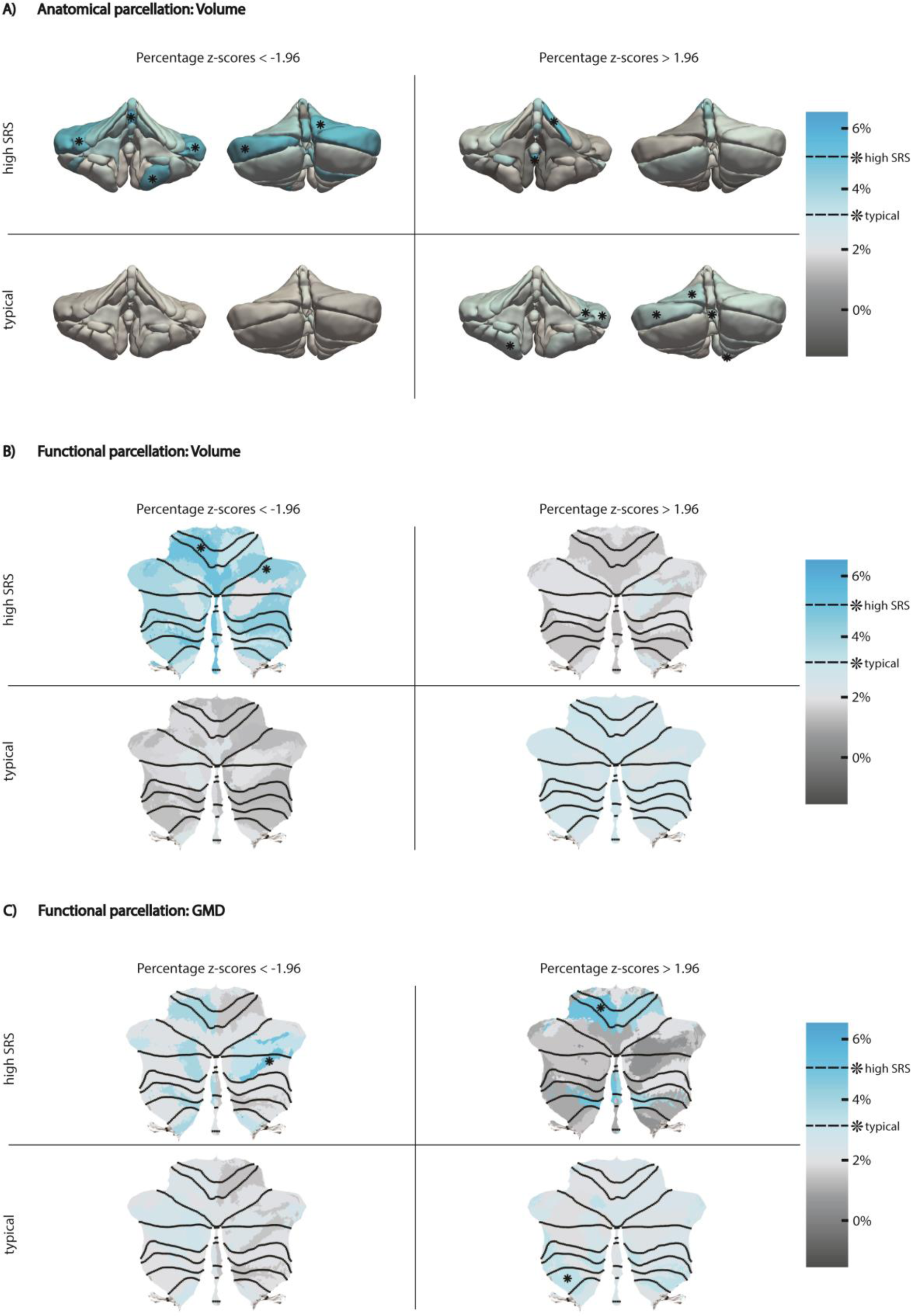
Percentage of individuals with large negative (z-score < −1.96) and large positive (z-score > 1.96) deviations in typically developing children and children that are likely to fall on the autism spectrum (high SRS score). Asterisks indicate ROIs in which children with high SRS and children with typical SRS scores have a significantly higher percentage of large deviation than expected (typical > 3.13%, high SRS > 5.05%; binomial test, *p*<0.05). (**A)** Deviations in volume in the anatomical ROIs. **B)** Deviations in volume in functional ROIs. **C)** Deviations in *Grey Matter Density* (GMD) in functional ROIs. *White Matter Density* (WMD) deviations are shown in *Supplementary Figure 4*.

Analogous to the anatomical parcellation, we again observe smaller volumes than expected throughout almost all functional parcels in participants with high SRS scores (**Figure 5B**). But interestingly, subtle distinctions are revealed in the functional parcellation. While an overall trend towards smaller volumes than expected is visible in participants with high SRS scores (significant percentage with large deviations binomial test at p<0.05: 1) *Left-hand presses* and 6) *Divided attention (right)*), the functional parcels best characterized by the cognitive features 7) *narrative* and 8) *word comprehension* seem to be exempted from this. Looking at the GMD maps of large deviations in high SRS participants, more negative deviations than expected (lower GMD) in the left anterior cerebellum and posterior cerebellum, specifically in parcellation best characterized by cognitive feature 9) *verbal fluency* (*p*<0.05), can be observed. Moreover, we clearly see more positive deviations than expected (higher GMD) in the anterior motor areas of the functional atlas (1) *Left-hand presses* (*p*<0.05) and 2) *Right-hand presses* in high SRS participants, as well as significantly higher GMD than expected in the functional region 4) *Action observation* (*p*<0.05) in typical participants (**Figure 4C**). Less pronounced differences are apparent in WMD deviations (*Supplementary Figure 4*).

## 5. Discussion

This study describes normative models of typical cerebellar development in a large pediatric population. Using over 7,000 longitudinal MRI scans, normative estimates of cerebellar growth from both anatomical and functional parcels were obtained by fitting hierarchical Bayesian regression models. Despite its potential to serve as a biomarker for developmental disorders (Fatemi et al., 2012; Sathyanesan et al., 2019; Wang et al., 2014), the morphology of the cerebellum has never been studied at this scale and detail using human MRI data before. Previous projects on normative modeling have demonstrated the potential impact and demand for large, open-source growth models (Bethlehem et al., 2022; Rutherford et al., 2022a). However, presently available models fail to include the cerebellum. The current models will therefore be critical to understand the heterogeneity in cerebellar development as well as deviations from the normative range in disorders. Notably, these deviations can be investigated on the level of a single individual using the current approach.

The anatomical regions (MAGeT atlas, (Park et al., 2014)) and functional regions (MDTB atlas, (King et al., 2019)) show similar overall growth trends. As expected in this age range traversing late-childhood into adolescence, we see increasing volumes throughout all ROIs in both parcellations (**Figure 2A, Figure 4A I&IV**). Slight sex differences can be found in the posterior cerebellar GMD and WMD maps (**Figure 4A II-III&V-VI**), with females showing slower GMD decreases and WMD increases compared to males in the right cerebellar hemisphere, possibly indicating an approaching developmental ceiling in females. Prolonged growth trajectories in males have been reported previously in the cerebellum (Tiemeier et al., 2010) and can also be seen in the cerebrum (Lenroot et al., 2007).

Consistent with previous findings, we observed an anterior-posterior gradient in cerebellar development likely to reflect and mirror the age-related improvements in underlying functions, with sensorimotor areas predominantly located anteriorly and cognitive areas posteriorly in the cerebellum (King et al., 2019; Klein et al., 2016; Liu et al., 2022). Anterior sensorimotor areas show smaller age-related effects compared to posterior cognitive areas, possibly reflecting protracted growth trajectories for higher-order cognitive compared to sensorimotor regions in the cerebellum (**Figure 2 & Figure 4**). This is evident in both males and females and throughout different morphological indicators (i.e., volumes as well as GMD, and WMD) but better captured by the functional than the anatomical parcellation. As functional activity in the brain rarely coincides with macroscopic anatomy (Brett et al., 2002) future studies employing functional parcellations might be able to uncover predictors of behavior in clinical subgroups that would otherwise be hidden using anatomical atlases only.

Interestingly, the reported developmental trends in the cerebellum closely mirror those found in the cerebrum. Similar patterns of earlier maturation of sensorimotor compared to higher order cognitive areas can be observed in myelination (Deoni et al., 2015; Elston & Fujita, 2014) and grey matter maturation (Giedd et al., 2015; Gilmore et al., 2012; Gogtay & Thompson, 2010; Tamnes et al., 2017) in the cerebral cortex, pointing towards directly related growth trajectories of the cerebellum and the cerebrum. Indeed, the cerebellum has been proposed as a crucial node for optimal structural and functional brain development. Hence, Wang and colleagues have recently coined the term of a *developmental diaschisis,* suggesting that the cerebellum might have a direct influence on cortical maturation (Wang et al., 2014). This also accords with earlier findings of volume decreases in remote but connected cerebral regions after perinatal cerebellar injury (Limperopoulos et al., 2014) and findings of cerebellar tumors resulting in significant downstream effects on higher cognition and motor function which could not be compensated well by other structures (Davis et al., 2010). Further work, in particular in vivo human research, is required to develop a more complete understanding of the cerebellar influence on cortical development.

An increasing amount of literature about the role of the cerebellum in higher cognitive functions but also in neurodevelopmental disorders, and in ASD specifically, has become available in recent years. Cerebellar abnormalities are among the most frequently reported in ASD patients (Wang et al., 2014). Intriguing reports from mouse models show that targeted activation of right Crus I and the posterior vermis was able to rescue autistic behaviors in TSC1 mutant mice by modulating activity in the medial prefrontal cortex (Kelly et al., 2020). Akin volumetric changes in Crus I and the posterior vermis have been reported in human studies as well as deviations in total cerebellar size (Courchesne et al., 2001; D’Mello et al., 2015; Kaufmann et al., 2003; McKinney et al., 2022; Pierce & Courchesne, 2001; Stanfield et al., 2008). Recently, however, no differences in cerebellar anatomy in individuals with autistic traits were reported when using normative models on cerebellar growth based on a smaller control sample (N=219) (Laidi et al., 2022). Therefore, looking into cerebellar deviations of children that are likely to fall on the Autism spectrum in large, population-based cohorts, lends itself as a prime example of illustrating the utility of the current normative models.

In accordance with previous research, we find smaller cerebellar volumes in children with autistic traits. In the anatomical parcellation smaller volumes can be seen throughout various regions, particularly in vermal and lobular parts of the anterior and superior posterior cerebellum (**Figure 5A**). While the functional parcellation also reveals smaller volumes throughout almost all ROIs, with significant differences in MDTB components 1) *Left-hand presses* and 6) *Divided attention (right)*, the MDTB components *7) Narrative* and *8) Word comprehension* seem to not follow the same trend (**Figure 5B**). Given the overlap of MDTB components 7 and 8 with the cerebellar default mode regions described by Buckner and colleagues (Buckner et al., 2011), a network found to be among the most disrupted in ASD patients, this might relate to previously reported heterogeneity in default mode network connectivity in children on the Autism spectrum (Padmanabhan et al., 2017). Clear differences can also be observed in GMD with a high percentage of individuals with autistic traits exhibiting increased GMD in the anterior sensorimotor parcels, particularly on the left hemisphere, and decreased GMD in the superior posterior parcels involved in language processing (**Figure 5C**). In view of the heterogeneity in brain morphology between individuals across a multitude of pathologies, it is noteworthy that the current approach does not depend on group level inferences but can be used at an individual level to uncover within-group heterogeneity.

While we chose to illustrate cerebellar deviations using the example of autistic traits, it is important to note that the cerebellum is known to play an influential role in a myriad of clinical subpopulations for which this approach would be particularly insightful. Following the concept of the *cerebellar connectome*, a framework proposing that deviations in cerebellar and cerebello-cortical connectivity have a direct influence on onset and severity of neurodevelopmental disorders, the current approach has the potential to not only advance our understanding of disease etiology, but might also uncover new sites for therapeutic interventions (Sathyanesan et al., 2019).

Normative models of all ROIs described are freely available on the PCNportal (https://pcnportal.dccn.nl/). Both linear as well as b-spline models can be downloaded and used as informative priors for new unseen sites using the PCNtoolkit (https://github.com/amarquand/PCNtoolkit.git). This not only allows for better predictions in smaller data sets, but it also enables future studies to model individual differences free of site-effects, and able to uncover clinically significant deviations from the normative model. To transfer knowledge from the current models to a new cohort an adaptation set of approximately 25 samples is needed (Gaiser et al., 2023). The remainder of the cohort can then be interpreted on a single subject basis and compared with the reference model without the need to employ an additional control group. A detailed account of how normative models can be implemented in future studies can be found online (https://pcntoolkit.readthedocs.io/en/latest/) and is also described by Gaiser, Berthet and colleagues (Gaiser et al., 2023). The PCNportal further offers the possibility to derive subject-level statistics in a new dataset in a simple and accessible way, without the need for any technical background knowledge or computing power. Greater efforts are needed to discern the role of the cerebellum in typical and atypical neurodevelopment for which the current normative models can serve as a highly useful tool.

### 5.1. Limitations

The Generation R Study is a population-wide cohort from The Netherlands and therefore the current normative models might not generalize ideally to other populations. As is common in brain imaging research (Henrich et al., 2010), families with high *socioeconomic status* (SES) and children with above average IQ are overrepresented in this cohort relative to the original catchment area. Fortunately, the current models can easily be updated within the PCNtoolkit framework by including new data points outside of our age range and from more diverse datasets, including different populations and clinical phenotypes. As a consequence, the cerebellar normative models can be extended and refined as new information becomes available while new, possibly smaller cohorts can benefit from informed priors based on our models. Furthermore, our scans were acquired on 3T MRI scanners with a voxel resolution of 1 mm^3^. While this standard setup is able to reliably identify cerebellar lobules (Park et al., 2014), higher resolutions are needed to more accurately segment cerebellar vermal regions, and white and grey matter given the thin, tightly folded cortical layering of the cerebellar cortex (Marques et al., 2010). Lastly, the functional parcellation used in this study was a group-average parcellation derived from high-quality, extensive functional MRI assessment from adults (King et al., 2019). Functional boundaries are likely to vary between individuals to some degree and may also vary as a function of age. Future studies should therefore aim to repeat the task-battery in a cohort of young children and adolescents to quantify whether neurodevelopmental differences exist in the functional parcellation.

## 6. Conclusion

In the current study, we present models of cerebellar growth during childhood and adolescence, an important time period for brain development, based on a large, prospective population cohort, the Generation R study. We find an anterior-posterior growth gradient mirroring the age-related improvements of underlying behavior and function. The anterior/sensorimotor-posterior/cognitive growth gradient mirrors cerebral maturation patterns, thus providing evidence for directly related cerebello-cortical developmental trajectories. In recent years, the cerebellum has received increasing attention as a critical node in fundamental cognitive and emotional functions as well as brain development. The current openly accessible growth models will therefore be of great value for uncovering cerebellar deviations and understanding their implications in neuropathology.

## Acknowledgments

The authors thank Nadine Danner for support in quality control of the MRI images, Jonathan Krikeb for help with data management and Min Tae M. Park for assisting with implementation of the MAGeT anatomical segmentation algorithm. Importantly, we thank and are grateful for the contribution of children and parents of the Generation R study, as well as the researchers involved in data collection.

## Data Availability

The cerebellar growth models are available on the PCNportal (https://pcnportal.dccn.nl/). MRI data from the Generation R cohort are not publicly available due to legal and ethical restrictions, however access can be requested via the Generation R administration.

## Code Availability

Code to generate normative models and transfer knowledge from existing models to new sites is freely available via the PCNtoolkit (https://github.com/amarquand/PCNtoolkit). Further codes are available from the corresponding author upon reasonable request.

