## Supplementary material for "Large Data on the Small Brain: Population-wide Cerebellar Growth Models of Children and Adolescents"

### Supplementary Figures

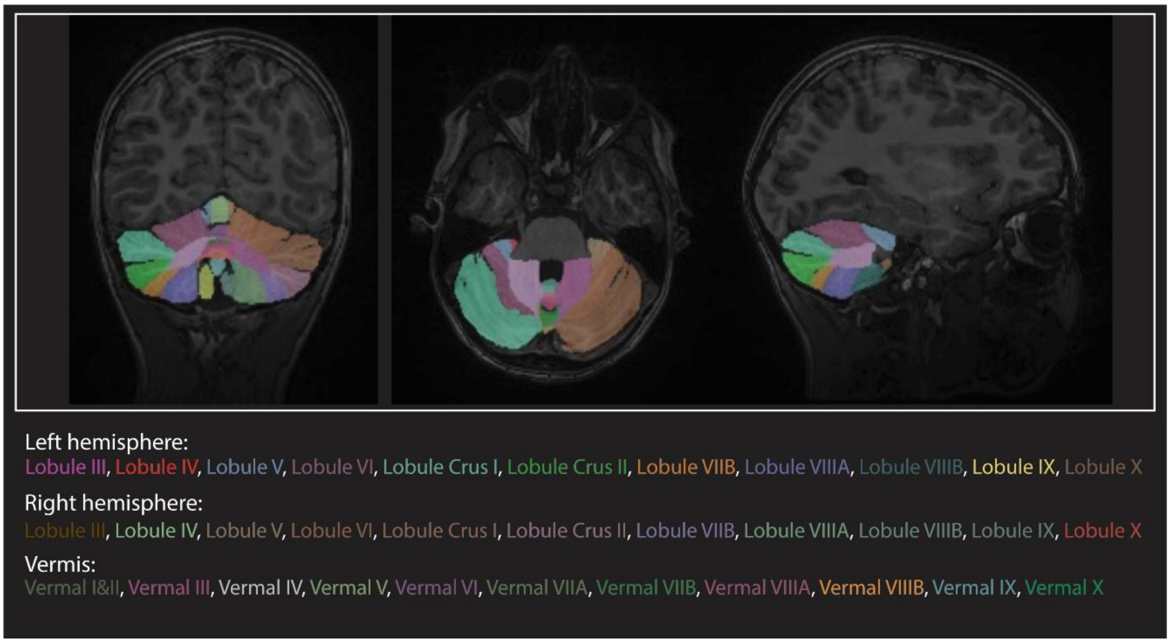


**Supplementary Figure 1**: Example scan that was automatically segmented in anatomical ROIs using the MAGeT algorithm. Labels for each ROI in their respective colors are shown.

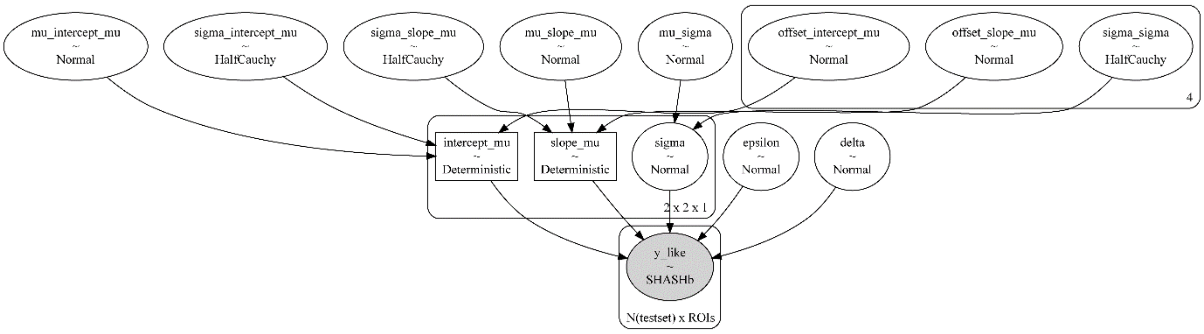


**Supplementary Figure 2**: Graphical representation of the normative model. Hierarchical structure, model parameters and their relationships between parameters are shown. Batch-effects are shown as 2x2 design (2 scanners x 2 sexes).

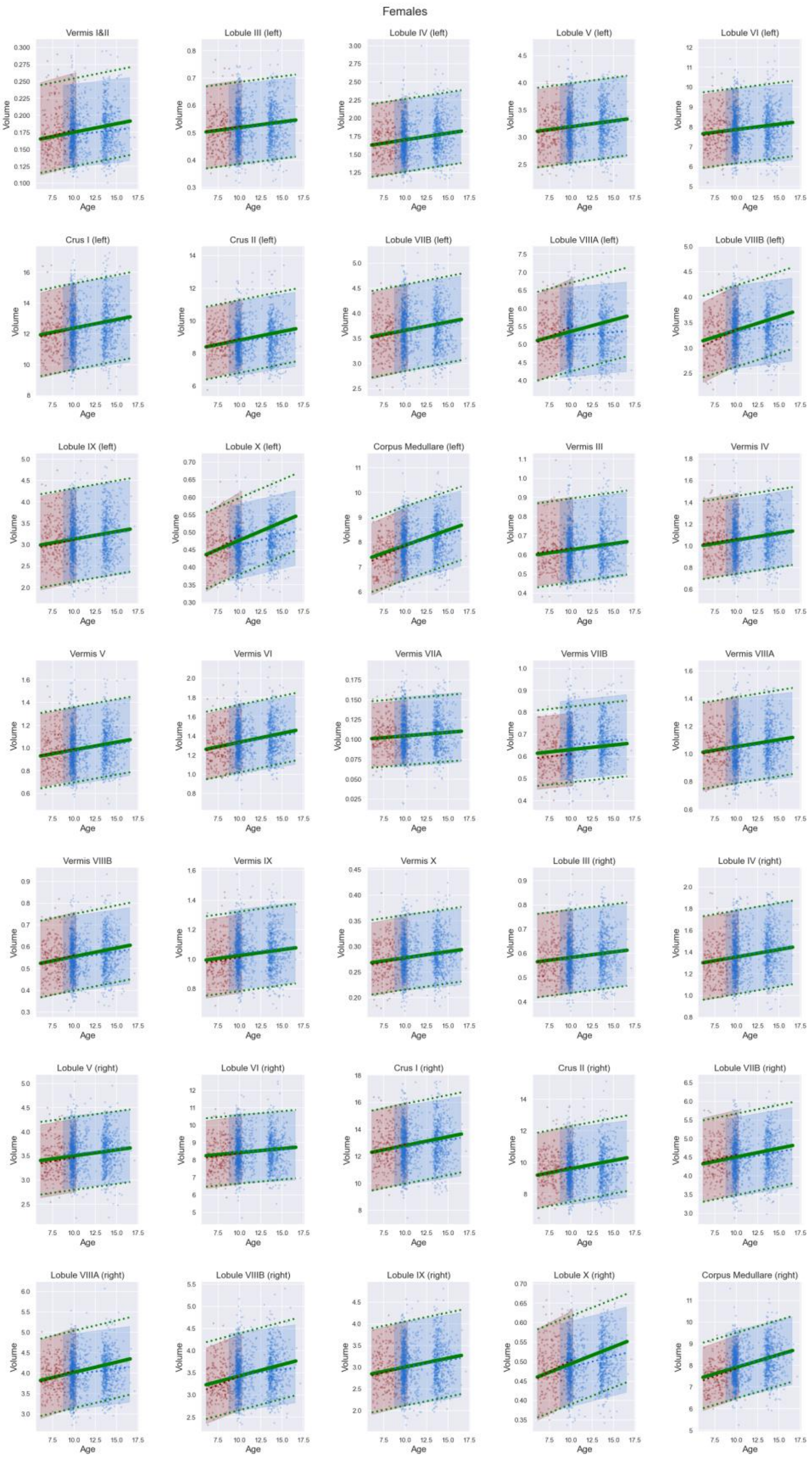


**Supplementary Figure 3A**: Growth trajectory for each anatomical ROI for all females. Bold green lines show the mean trajectory, dotted green lines represent what is within 2 standard deviations of the mean. In red all data points of females acquired on the first scanner (visit/wave 1) are shown. Red dotted line and red shaded area illustrate the mean trajectory and what is within 2 standard deviations considering the batch-effect of the first scanner only. Analogous in blue, data points of females acquired on the second scanner (visit/wave 2 & 3) are shown. Blue dotted line and blue shaded area illustrate the mean trajectory and what is within 2 standard deviations considering the batch-effect of the second scanner only.

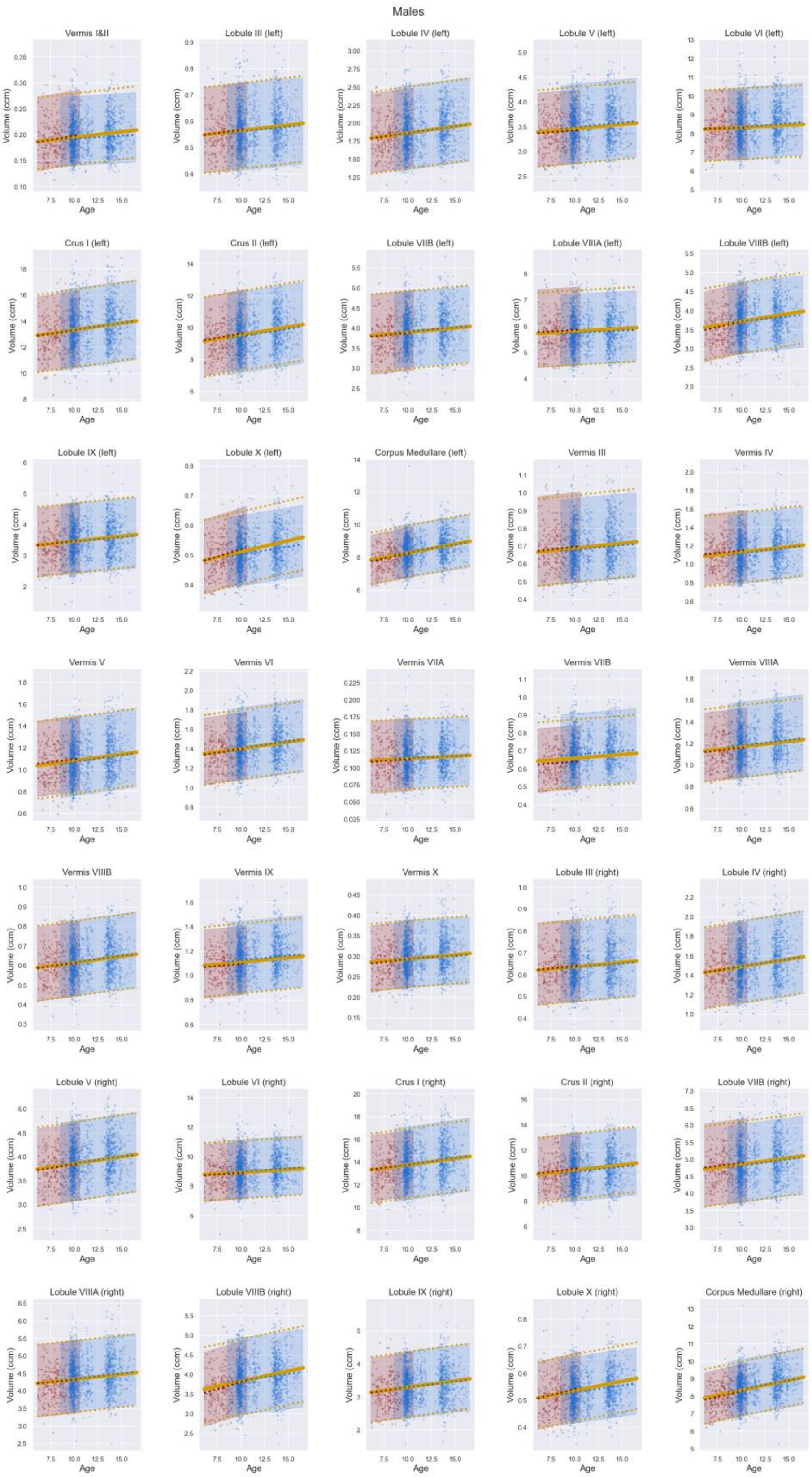


**Supplementary Figure 3B**: Growth trajectory for each anatomical ROI for all males. Bold yellow lines show the mean trajectory, dotted yellow lines represent what is within 2 standard deviations of the mean. In red all data points of males acquired on the first scanner (visit/wave 1) are shown. Red dotted line and red shaded area illustrate the mean trajectory and what is within 2 standard deviations considering the batch-effect of the first scanner only. Analogous in blue, data points of males acquired on the second scanner (visit/wave 2 & 3) are shown. Blue dotted line and blue shaded area illustrate the mean trajectory and what is within 2 standard deviations considering the batch-effect of the second scanner only.

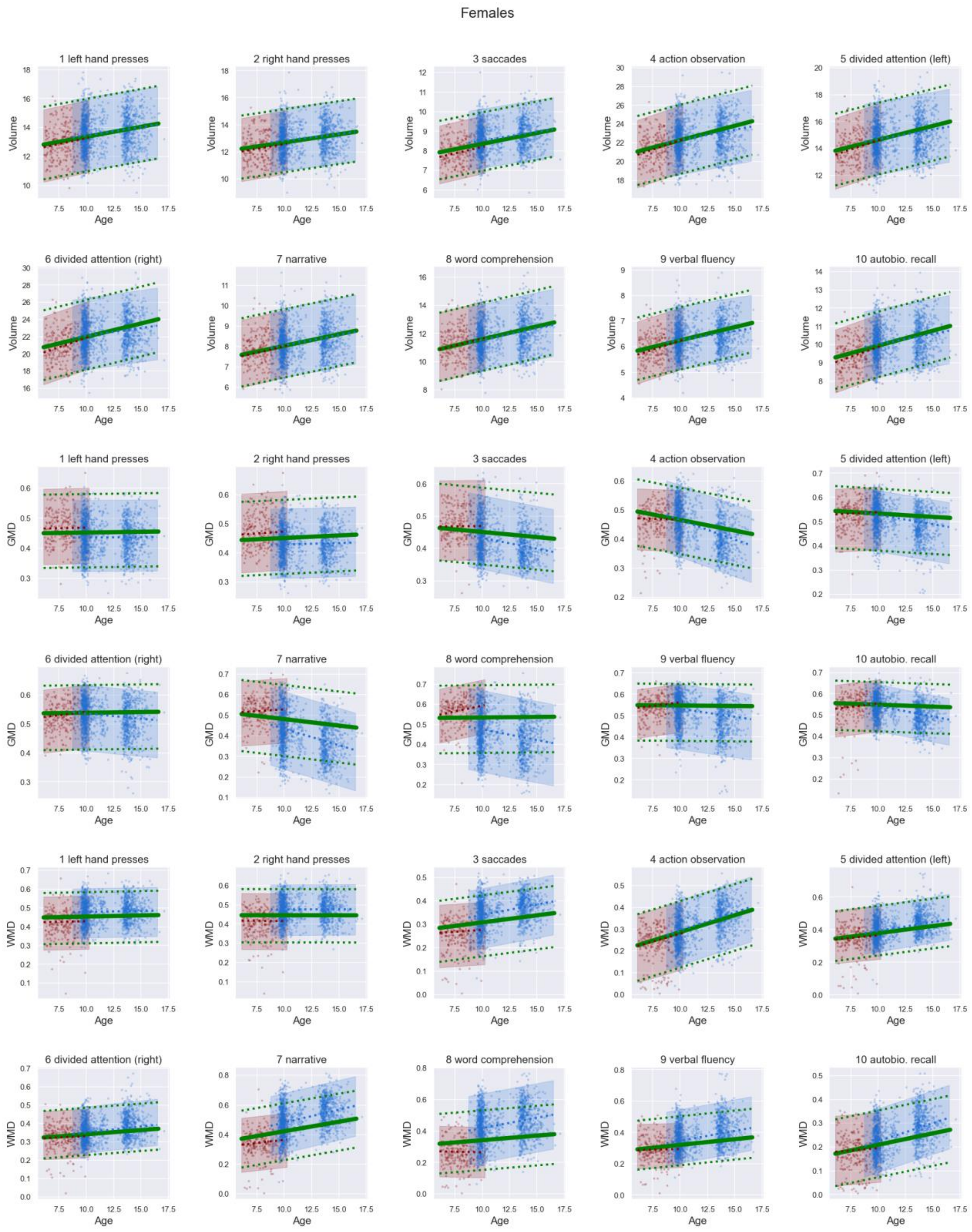


**Supplementary Figure 3C:** Growth trajectory for each functional ROI for all females. First 2 rows depict trajectories for volumes, 3rd and 4th row for GMD, 5th and 6th row for WMD. Bold green lines show the mean trajectory, dotted green lines represent what is within 2 standard deviations of the mean. In red all data points of females acquired on the first scanner (visit/wave 1) are shown. Red dotted line and red shaded area illustrate the mean trajectory and what is within 2 standard deviations considering the batch-effect of the first scanner only. Analogous in blue, data points of females acquired on the second scanner (visit/wave 2 & 3) are shown. Blue dotted line and blue shaded area illustrate the mean trajectory and what is within 2 standard deviations considering the batch-effect of the second scanner only.

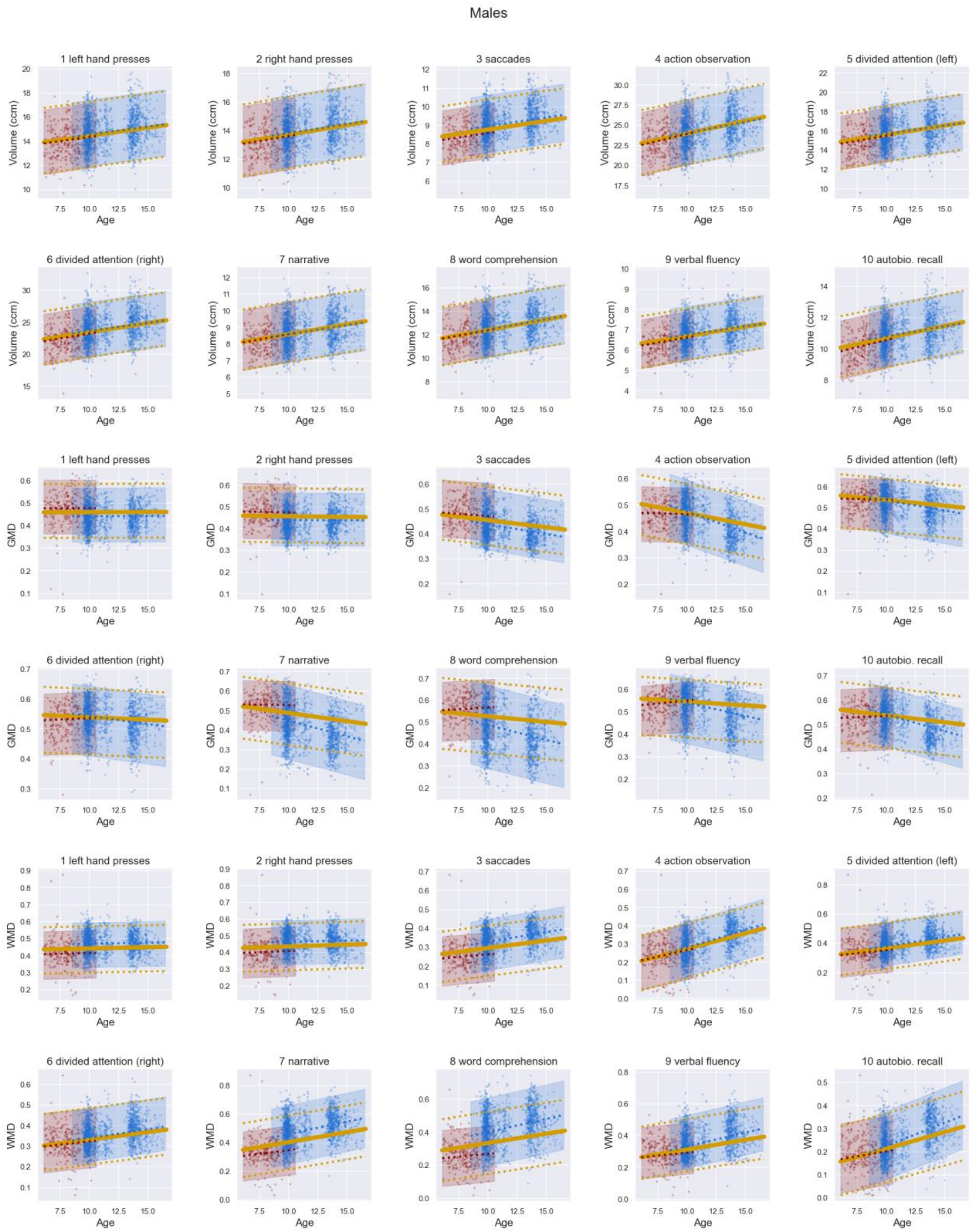


**Supplementary Figure 3D**: Growth trajectory for each functional ROI for all males. First 2 rows depict trajectories for volumes, 3rd and 4th row for GMD, 5th and 6th row for WMD. Bold yellow lines show the mean trajectory, dotted yellow lines represent what is within 2 standard deviations of the mean. In red all data points of males acquired on the first scanner (visit/wave 1) are shown. Red dotted line and red shaded area illustrate the mean trajectory and what is within 2 standard deviations considering the batch-effect of the first scanner only. Analogous in blue, data points of males acquired on the second scanner (visit/wave 2 & 3) are shown. Blue dotted line and blue shaded area illustrate the mean trajectory and what is within 2 standard deviations considering the batch-effect of the second scanner only.

**Functional parcellation: WMD**

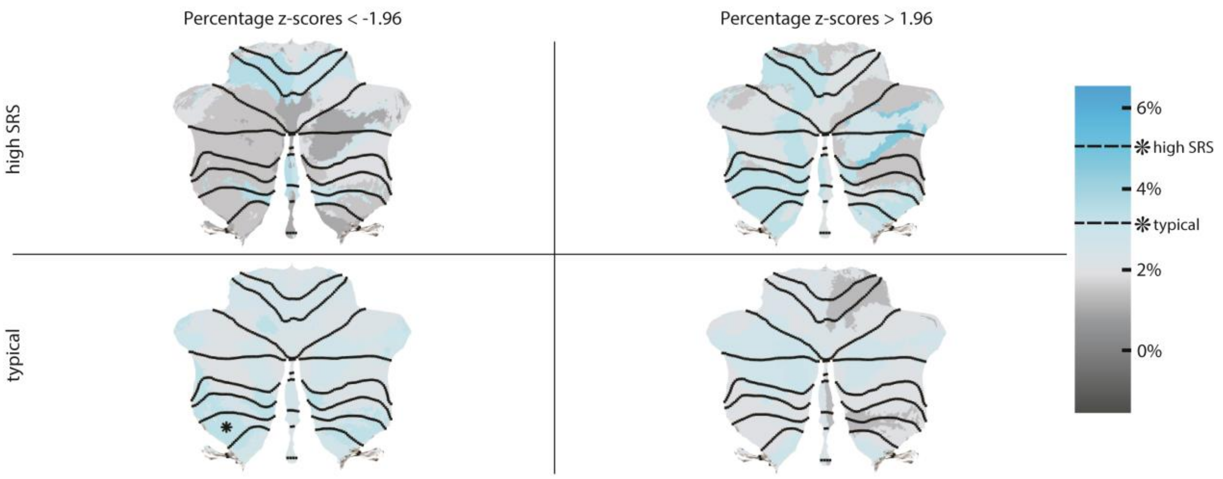


**Supplementary Figure 4**: Percentage of individuals with large negative (z-score < -1.96) and large positive (z-score > 1.96) deviations in *White Matter Density* (WMD) in functional ROIs. Asterisks indicate ROIs in which children with high SRS and children with typical SRS scores have a significantly higher percentage of large deviation than expected (typical > 3.13%, high SRS > 5.05%; binomial test, $p$<0.05). **Top row**: children with high SRS scores, likely to fall on the Autism spectrum. **Bottom row**: typically developing children.

### Supplementary Tables

**Supplementary Table 1:** *sample characteristics of the training and test sets*

<table>
<tr><th rowspan="2"></th><th colspan="3">Training set<br>N = 3,689</th><th colspan="3">Test set<br>N = 3,581</th><th rowspan="2">Total<br>N = 7,270</th></tr>
<tr><th>Wave 1<br>N = 496</th><th>Wave 2<br>N = 1,961</th><th>Wave 3<br>N = 1,319</th><th>Wave 1<br>N = 483</th><th>Wave 2<br>N = 1,968</th><th>Wave 3<br>N = 1,224</th></tr>
<tr><td>Age<br>Mean [Range]</td><td>7.9 [6.1 – 10.7]</td><td>10.1 [8.7 – 12.0]</td><td>14.0 [12.7 – 16.6]</td><td>7.9 [6.2 – 10.7]</td><td>10.1 [8.6 – 12.0]</td><td>14.0 [12.6 – 17.1]</td><td>11.2 [6.1 – 17.1]</td></tr>
<tr><td>Sex<br>(M|F)</td><td>53.8% | 46.2%</td><td>48.1% | 51.9%</td><td>47.6% | 52.4%</td><td>50.9% | 49.1%</td><td>51.2% | 48.8%</td><td>48.2% | 51.8%</td><td>49.4% | 50.6%</td></tr>
<tr><td>Income<br>(Low | Medium | High)</td><td>4.9% | 38.3% | 47.9%</td><td>6.0% | 31.2% | 49.8%</td><td>4.5% | 34.3% | 49.7%</td><td>7.7% | 38.9% | 45.9%</td><td>4.8% | 34.1% | 49.1%</td><td>5.0% | 33.2% | 47.7%</td><td>5.3% | 33.8% | 48.9%</td></tr>
<tr><td>Education Mother<br>(Low | High)</td><td>39.6% | 60.4%</td><td>33.5% | 66.5%</td><td>34.1% | 65.9%</td><td>42.2% | 57.8%</td><td>34.9% | 65.1%</td><td>35.6% | 64.4%</td><td>35.3% | 64.7%</td></tr>
<tr><td>IQ<br>(Low | Medium | High)</td><td>12.2% | 59.8% | 15.6%</td><td>9.1% | 56.9% | 15.8%</td><td>8.5% | 57.7% | 17.2%</td><td>11.4% | 60.3% | 16.2%</td><td>10.1% | 53.7% | 18.3%</td><td>10.6% | 55.7% | 15.5%</td><td>9.8% | 56.4% | 16.7%</td></tr>
<tr><td>Ethnicity<br>(Dutch | Western (Non-Dutch) | Non-Western)</td><td>68.0% | 6.5% | 25.6%</td><td>59.6% | 8.5% | 30.4%</td><td>60.0% | 8.3% | 29.5%</td><td>67.8% | 6.7% | 25.6%</td><td>58.2% | 8.4% | 30.9%</td><td>57.6% | 9.2% | 30.9%</td><td>60.1% | 8.3% | 29.8%</td></tr>
<tr><td>CBCL<br>(Low | High )</td><td>58.6% | 22.9%</td><td>67.1% | 17.1%</td><td>64.3% | 17.0%</td><td>57.6% | 19.3%</td><td>67.7% | 16.9%</td><td>65.0% | 14.3%</td><td>65.2% | 17.1%</td></tr>
</table>

** percentages do not always add up to 100% due to missing data from some participants*

| ROI | linear<br>loo [SE] | bspline<br>loo [SE] | difference [SE of difference] |
|---|---|---|---|
| Vermis I & II | -4910.89 [45.54] | -4911.74 [45.45] | 0.85 [2.58] |
| Lobule III (left) | -5034.17 [43.25] | -5036.97 [43.29] | 2.79 [2.51] |
| Lobule IV (left) | -4966.64 [44.45] | -4984.73 [44.56] | 2.10 [2.45] |
| Lobule V (left) | -4940.60 [42.93] | -4945.15 [43.04] | 4.55 [1.95] |
| Lobule VI (left) | -5078.43 [45.37] | -5082.04 [45.35] | 3.61 [1.73] |
| Lobule Crus 1 (left) | -4992.30 [43.99] | -4995.34 [43.96] | 3.04 [1.67] |
| Lobule Crus 2 (left) | -4981.02 [45.78] | -4980.35 [45.78] | 0.67 [3.10] |
| Lobule VIIB (left) | -5066.31 [44.01] | -5069.97 [44.05] | 3.66 [1.88] |
| Lobule VIIIA (left) | -4915.24 [45.22] | -4914.21 [45.11] | 1.03 [3.25] |
| Lobule VIIIB (left) | -4800.77 [45.32] | -4799.91 [45.44] | 0.86 [3.30] |
| Lobule IX (left) | -5040.51 [46.02] | -5042.22 [46.03] | 1.72 [2.75] |
| Lobule X (left) | -4914.25 [45.01] | -4913.38 [44.96] | 0.87 [3.47] |
| Corpus medullare (left) | -4929.63 [45.65] | -4925.68 [45.68] | 3.95 [3.79] |
| Vermis III | -4961.70 [46.54] | -4965.54 [46.48] | 3.84 [1.89] |
| Vermis IV | -5067.51 [45.57] | -5072.15 [45.20] | 4.64 [1.43] |
| Vermis V | -5005.63 [42.50] | -5009.64 [42.44] | 4.01 [1.68] |
| Vermis VI | -5060.31 [44.72] | -5063.01 [44.90] | 2.71 [2.31] |
| Vermis VIIA | -5084.65 [47.27] | -5089.64 [47.14] | 4.99 [1.63] |
| Vermis VIIB | -5050.23 [44.62] | -5056.92 [44.61] | 6.70 [1.10] |
| Vermis VIIIA | -4840.80 [45.91] | -4843.85 [45.90] | 3.06 [2.24] |
| Vermis VIIIB | -4916.88 [43.85] | -4920.07 [43.98] | 3.19 [2.13] |
| Vermis IX | -4994.13 [45.09] | -4997.92 [45.06] | 3.79 [2.15] |
| Vermis X | -5056.58 [48.49] | -5058.67 [48.53] | 2.10 [2.17] |
| Lobule III (right) | -5017.72 [45.46] | -5020.44 [45.58] | 2.72 [2.00] |
| Lobule IV (right) | -4966.79 [43.61] | -4968.72 [43.56] | 1.93 [2.13] |
| Lobule V (right) | -4835.30 [45.11] | -4838.61 [45.17] | 3.31 [1.99] |
| Lobule VI (right) | -5067.93 [45.63] | -5072.41 [45.72] | 4.48 [1.41] |
| Lobule Crus 1 (right) | -4974.52 [43.50] | -4977.94 [43.50] | 3.42 [1.90] |
| Lobule Crus 2 (right) | -4970.96 [44.83] | -4972.03 [44.79] | 1.07 [2.73] |
| Lobule VIIB (right) | -4997.19 [43.77] | -5001.44 [43.69] | 4.25 [2.10] |
| Lobule VIIIA (right) | -4920.04 [45.58] | -4921.01 [45.61] | 0.97 [2.76] |
| Lobule VIIIB (right) | -4778.78 [45.31] | -4781.65 [45.30] | 2.87 [2.44] |
| Lobule IX (right) | -5018.76 [44.65] | -5018.51 [44.57] | 0.25 [3.17] |
| Lobule X (right) | -4905.17 [43.62] | -4908.13 [43.62] | 2.96 [2.34] |
| Corpus medullare (right) | -4892.49 [46.78] | -4892.19 [46.79] | 0.30 [3.29] |
| left hand presses (Volume) | -4814.67 [44.80] | -4818.54 [44.91] | 3.87 [2.11] |
| right hand presses (Volume) | -4783.62 [45.92] | -4786.92 [46.00] | 3.30 [2.13] |
| saccades (Volume) | -4822.83 [44.19] | -4825.58 [44.40] | 2.75 [2.40] |
| action observation (Volume) | -4708.00 [44.40] | -4709.21 [44.45] | 1.22 [2.73] |
| divided attention (left) (Volume) | -4838.92 [44.82] | -4838.44 [44.80] | 0.47 [3.15] |
| divided attention (right) (Volume) | -4808.71 [46.02] | -4811.82 [46.02] | 3.11 [2.00] |
| narrative (Volume) | -4924.14 [42.77] | -4923.69 [42.69] | 0.45 [3.35] |
| word comprehension (Volume) | -4833.15 [43.01] | -4834.34 [43.07] | 1.19 [2.62] |
| verbal fluency (Volume) | -4776.38 [44.56] | -4778.28 [44.60] | 1.91 [2.68] |
| autobiographical recall (Volume) | -4728.35 [44.27] | -4729.09 [44.36] | 0.74 [2.80] |
| left hand presses (GMD) | -5166.46 [44.01] | -5156.58 [44.39] | 9.87 [5.50] |
| right hand presses (GMD) | -5147.05 [45.60] | -5142.16 [45.83] | 4.89 [4.03] |
| saccades (GMD) | -4930.29 [48.30] | -4920.94 [48.40] | 9.34 [4.39] |
| action observation (GMD) | -4857.93 [43.38] | -4845.04 [43.32] | 12.89 [6.43] |
| divided attention (left) (GMD) | -4998.79 [46.76] | -4989.39 [46.91] | 9.40 [5.10] |
| divided attention (right) (GMD) | -5135.83 [45.19] | -5130.88 [45.38] | 4.95 [4.15] |
| narrative (GMD) | -4743.98 [45.23] | -4729.98 [45.35] | 14.00 [5.59] |
| word comprehension (GMD) | -4840.82 [48.20] | -4832.78 [48.15] | 8.05 [5.07] |
| verbal fluency (GMD) | -4882.19 [46.91] | -4876.16 [47.06] | 6.03 [5.09] |
| autobiographical recall (GMD) | -4939.08 [48.31] | -4916.27 [48.08] | 22.81 [7.89] |
| left hand presses (WMD) | -5067.61 [47.37] | -5055.36 [47.57] | 12.25 [5.83] |
| right hand presses (WMD) | -5037.57 [48.02] | -5031.54 [48.34] | 6.02 [4.45] |
| saccades (WMD) | -4717.58 [50.32] | -4702.22 [50.21] | 15.35 [5.66] |
| action observation (WMD) | -4707.40 [43.81] | -4695.48 [44.00] | 11.92 [5.74] |
| divided attention (left) (WMD) | -4858.86 [49.10] | -4844.21 [49.28] | 14.65 [5.85] |
| divided attention (right) (WMD) | -5008.70 [47.83] | -5000.85 [48.34] | 7.85 [4.94] |
| narrative (WMD) | -4556.83 [46.12] | -4541.50 [46.25] | 15.33 [5.74] |
| word comprehension (WMD) | -4604.72 [48.00] | -4595.59 [47.95] | 9.14 [5.11] |
| verbal fluency (WMD) | -4756.39 [46.60] | -4744.21 [46.94] | 12.18 [5.85] |
| autobiographical recall (WMD) | -4696.75 [45.18] | -4665.38 [45.19] | 31.37 [8.81] |

**Supplementary Table 2**: Leave-one-out (LOO) Cross Validation. LOO, standard error (SE) for the LOO computations of linear and b-spline models are shown as well for difference in LOO and the SE of the difference for all anatomical and functional ROIs.

| Anatomical lobular ROIS | **Volume**<br>**Mean standardized β Age [95%CI Mean]**<br>*Left hemisphere* | *Right hemisphere* | Anatomical vermal ROIs | **Volume**<br>**Mean standardized β Age [95%CI Mean]** |
|---|---|---|---|---|
| | | | Vermis I&II | |
| | | | **MALES** | 0.142 [1.140 – 1.44] |
| | | | **FEMALES** | 0.167 [0.165 – 0.169] |
| Lobule III | | | Vermis III | |
| **MALES** | 0.116 [0.115 – 0.117] | 0.100 [0.099 - 0.100] | **MALES** | 0.107 [0.106 - 0.108] |
| **FEMALES** | 0.115 [0.114 – 0.116] | 0.112 [0.111 - 0.113] | **FEMALES** | 0.124 [0.123 - 0.124] |
| Lobule IV | | | Vermis IV | |
| **MALES** | 0.148 [0.147 – 0.149] | 0.162 [0.162 - 0.163] | **MALES** | 0.138 [0.137 - 0.139] |
| **FEMALES** | 0.146 [0.146 – 0.147] | 0.144 [0.143 - 0.145] | **FEMALES** | 0.151 [0.150 - 0.152] |
| Lobule V | | | Vermis V | |
| **MALES** | 0.095 [0.094 – 0.096] | 0.154 [0.153 - 0.155] | **MALES** | 0.144 [0.143 - 0.145] |
| **FEMALES** | 0.117 [0.116 – 0.118] | 0.130 [0.129 - 0.132] | **FEMALES** | 0.169 [0.168 - 0.170] |
| Lobule VI | | | Vermis VI | |
| **MALES** | 0.058 [0.056 – 0.059] | 0.086 [0.085 - 0.087] | **MALES** | 0.165 [0.164 - 0.166] |
| **FEMALES** | 0.126 [0.124 – 0.128] | 0.102 [0.101 - 0.104] | **FEMALES** | 0.226 [0.224 - 0.228] |
| Crus I | | | Vermis VIIA | |
| **MALES** | 0.153 [0.152 – 0.154] | 0.156 [0.155 - 0.157] | **MALES** | 0.069 [0.068 - 0.070] |
| **FEMALES** | 0.161 [0.160 – 0.162] | 0.179 [0.177 - 0.180] | **FEMALES** | 0.085 [0.084 - 0.086] |
| Crus II | | | | |
| **MALES** | 0.178 [0.177 – 0.179] | 0.145 [0.144 - 0.146] | | |
| **FEMALES** | 0.191 [0.189 – 0.192] | 0.179 [0.177 - 0.180] | | |
| Lobule VIIB | | | Vermis VIIB | |
| **MALES** | 0.104 [0.103 – 0.105] | 0.142 [0.141 - 0.143] | **MALES** | 0.098 [0.097 - 0.098] |
| **FEMALES** | 0.160 [0.158 – 0.161] | 0.181 [0.179 - 0.182] | **FEMALES** | 0.098 [0.097 - 0.098] |
| Lobule VIIIA | | | Vermis VIIIA | |
| **MALES** | 0.069 [0.067 – 0.070] | 0.126 [0.125 - 0.128] | **MALES** | 0.129 [0.128 - 0.130] |
| **FEMALES** | 0.210 [0.207 – 0.213] | 0.219 [0.217 - 0.222] | **FEMALES** | 0.130 [0.129 - 0.132] |
| Lobule VIIIB | | | Vermis VIIIB | |
| **MALES** | 0.192 [0.190 – 0.194] | 0.229 [0.227 - 0.231] | **MALES** | 0.149 [0.147 - 0.150] |
| **FEMALES** | 0.252 [0.249 – 0.255] | 0.227 [0.224 - 0.229] | **FEMALES** | 0.181 [0.180 - 0.183] |
| Lobule IX | | | Vermis IX | |
| **MALES** | 0.129 [0.127 - 0.130] | 0.171 [0.170 - 0.172] | **MALES** | 0.113 [0.112 - 0.114] |
| **FEMALES** | 0.141 [0.139 - 0.142] | 0.179 [0.177 - 0.180] | **FEMALES** | 0.121 [0.121 - 0.122] |
| Lobule X | | | Vermis X | |
| **MALES** | 0.280 [0.277 - 0.282] | 0.250 [0.248 - 0.251] | **MALES** | 0.115 [0.113 - 0.116] |
| **FEMALES** | 0.390 [0.386 - 0.394] | 0.312 [0.310 - 0.315] | **FEMALES** | 0.140 [0.139 - 0.142] |
| Corpus medullare | | | | |
| **MALES** | 0.293 [0.292 -0.295] | 0.303 [0.301 - 0.304] | | |
| **FEMALES** | 0.334 [0.331 - 0.336] | 0.315 [0.313 - 0.317] | | |

**Supplementary Table 3**: Mean standardized age β coefficients (slopes) and 95% confidence interval (CI) of the mean for all anatomical ROIs stratified by sex.

| FUNCTIONAL REGIONS | VOLUME<br><br>MEAN STANDARDIZED Β AGE [95%CI MEAN] | GMD<br><br>MEAN STANDARDIZED Β AGE [95%CI MEAN] | WMD<br><br>MEAN STANDARDIZED Β AGE [95%CI MEAN] |
|---|---|---|---|
| **1 LEFT-HAND PRESSES** | | | |
| Males | 0.211 [0.210 – 0.212] | 0.005 [0.005 - 0.006] | 0.047 [0.046 - 0.048] |
| Females | 0.217 [0.216 – 0.218] | 0.020 [0.019 - 0.021] | 0.039 [0.038 - 0.040] |
| **2 RIGHT-HAND PRESSES** | | | |
| Males | 0.227 [0.226 – 0.228] | -0.023 [-0.025 - -0.022] | 0.071 [0.070 - 0.073] |
| Females | 0.204 [0.203 – 0.205] | 0.065 [0.063 -0.066] | -0.002 [-0.003 - -0.001] |
| **3 SACCADES** | | | |
| Males | 0.254 [0.253 - 0.255] | -0.206 [-0.208 - -0.204] | 0.247 [0.245 - 0.249] |
| Females | 0.299 [0.297 - 0.301] | -0.114 [-0.117 - -0.111] | 0.184 [0.182 - 0.187] |
| **4 ACTION OBSERVATION** | | | |
| Males | 0.317 [0.316 -0.318] | -0.292 [-0.297 - -0.288] | 0.452 [0.450 - 0.454] |
| Females | 0.314 [0.312 - 0.316] | -0.250 [-0.253 - -0.246] | 0.411 [0.409 - 0.412] |
| **5 DIVIDED ATTENTION (LEFT)** | | | |
| Males | 0.279 [0.277 - 0.280] | -0.186 [-0.189 - -0.183] | 0.304 [0.302 - 0.306] |
| Females | 0.304 [0.302 - 0.306] | -0.093 [-0.097 - -0.090] | 0.251 [0.249 - 0.253] |
| **6 DIVIDED ATTENTION (RIGHT)** | | | |
| Males | 0.278 [0.277 - 0.280] | -0.070 [-0.073 - -0.067] | 0.252 [0.250 - 0.253] |
| Females | 0.305 [0.302 - 0.308] | 0.019 [0.016 - 0.022] | 0.154 [0.152 - 0.156] |
| **7 NARRATIVE** | | | |
| Males | 0.282 [0.281 - 0.284] | -0.185 [-0.189 - -0.182] | 0.269 [0.267 - 0.270] |
| Females | 0.269 [0.268 - 0.271] | -0.137 [-0.141 - -0.133] | 0.248 [0.246 - 0.250] |
| **8 WORD COMPREHENSION** | | | |
| Males | 0.300 [0.299 - 0.301] | -0.113 [-0.116 - -0.109] | 0.210 [0.208 - 0.212] |
| Females | 0.301 [0.300 - 0.303] | 0.012 [0.008 - 0.016] | 0.107 [0.104 - 0.110] |
| **9 VERBAL FLUENCY** | | | |
| Males | 0.298 [0.297 - 0.300] | -0.099 [-0.104 - -0.095] | 0.299 [0.296 - 0.301] |
| Females | 0.331 [0.328 - 0.333] | -0.013 [-0.017 - -0.009] | 0.176 [0.174 - 0.179] |
| **10 AUTOBIOGRAPHICAL RECALL** | | | |
| Males | 0.326 [0.324 - 0.328] | -0.214 [-0.219 - -0.209] | 0.408 [0.404 - 0.411] |
| Females | 0.342 [0.340 - 0.345] | -0.067 [-0.073 - -0.062] | 0.270 [0.267 - 0.274] |

**Supplementary Table 4:** Mean standardized age β coefficients (slopes) and 95% confidence interval (CI) of the mean for all functional ROIs stratified by sex.
